# Order code in the olfactory system

**DOI:** 10.1101/2025.11.16.688691

**Authors:** Khristina Samoilova, Joshua S. Harvey, Hirofumi Nakayama, Dmitry Rinberg, Alexei Koulakov

## Abstract

The ability to recognize odor identity across a wide range of concentrations is essential for natural olfactory behaviors. However, how odor identity is represented in the early olfactory system remains an open question. One theory proposes that glomeruli in the olfactory bulb, along with their associated odorant receptors (ORs), are activated by odorants in a temporal order that conveys information about odor identity. This order code is relatively robust to concentration changes as the rank order of glomerular activation by a given odorant remains similar across concentrations. Alternatively, the primacy coding theory suggests that the identity of each odorant is defined by its primacy set of glomeruli, comprised of the most sensitive ORs that respond first to that odorant. To test these theories, we measured glomerular responses to a large set of odorants in the mouse olfactory bulb using calcium imaging. We found that receptor affinities can be embedded in a low-dimensional space (D = 10) with minimal loss of information. Within this space, we identified two clusters of glomeruli with distinct tuning properties that form independent odor representations. These clusters may correspond to the two phylogenetic classes of ORs as revealed by both their functional characteristics and anatomical locations. In the OR affinity space, odorants evoke orderly activation waves whose directions can be used to define odor identities in the order coding model. We compared the order code and primacy model in predicting odor identity both across concentrations and across animals. Despite containing less information overall, the primacy model performed comparably to the order code. We confirmed the prediction of the primacy model that each receptor tends to contribute to the primacy set of at least one odorant. Analysis of binary odor mixtures revealed that mixture responses lie along geodesic lines connecting their component odors. Together, these findings suggest that odor information in the olfactory bulb may be represented through two complementary coding strategies: the primacy and order codes.

## INTRODUCTION

As with other sensory modalities, the olfactory system faces the problem of identifying a stimulus across variable contexts and concentrations. For example, the smell of food must signal consistent information at different distances to support successful navigation. The ability of the olfactory system to maintain a stable percept despite changes in odor concentration has been under intense study [1–8]. The question of odor identity stability can be framed as follows: which features of an odor’s neural representation remain consistent when the concentration of odorants varies? When identified, these features can be tested through direct experimental manipulation [6, 8–10] or by developing predictive computational models of odor identity [4, 8, 11, 12].

The olfactory system relies on odorant receptors (ORs) to detect and identify molecules in the air. ORs are specialized proteins that are expressed by the olfactory sensory neurons (OSNs) in the olfactory epithelium. Each OSN expresses one OR type from a repertoire of hundreds. OSNs expressing the same OR type project to discrete structures on the olfactory bulb’s (OB) surface, called glomeruli, with each glomerulus receiving inputs from a single OR type. Each odor activates a unique pattern of glomeruli, based on how different odorant receptors (ORs) respond to the molecule [13, 14]. These spatiotemporal activation patterns allow the olfactory system to encode odor identity and concentration. The challenge of finding the representation of odor identity is to discover the features of the neuronal activation patterns which are invariant with respect to odor concentration.

Several models have been proposed to explain how concentration-invariant odor identity is represented in the olfactory bulb (OB). Models based on receptor response magnitude can achieve concentration invariance by normalizing bulbar responses [15–18]. However, this approach only partially addresses the problem, because such normalization requires integrating inputs across all channels, or glomeruli [19], including those activated later in the sniff cycle [20]. This requirement seems to preclude odor-guided decisions based on early olfactory inputs [8, 21]. Temporal models of odor identity instead rely on the timing of glomerular/OR activation to represent odor identity. Hopfield suggested that the relative latencies of ORs activations could remain invariant to concentration for a given odor [22]. This assumption gave rise to the model of coincidence detectors in the olfactory (piriform) cortex that can detect the latencies to form an odor identity percept [23]. This model relies on the assumption that OR activation latencies depend on the logarithm of concentration and, therefore, the relative latencies are invariant to concentration changes, which may not be consistent with experimental findings [24, 25]. An alternative hypothesis states that odor identity is encoded not by the exact latencies of glomerular/OR activations, but by their relative order, the latency rank [26]. This model assumes that, despite variations in glomerular or OR latencies with concentration, their sorted rank order remains preserved. Consequently, Ref. [26] demonstrates that “order code” for odor identity yields a more robust odor classifier across trials than the latency model. Finally, a third model proposes that the earliest activated ORs or glomeruli, called the primacy set, carry the necessary information about odor identity. In this primacy model [8], most of the information contained in the OR activity patterns is discarded and odor identity emerges from distinguishing between early (primacy) and late (non-primacy) activated glomeruli or ORs. The primacy set represents a stable subset of the glomerular activity pattern, maintained across concentrations due to the consistent activation order of OR responses [8].

In this study, we performed wide-field 1-photon Ca imaging of glomerular responses to large panels of odors. We used glomerular activations to approximate the receptor affinities to odorants and organize the receptor odor space. The analysis of the receptor space revealed two independent channels which transmit odor information to the downstream networks. We then built representations of odor identity (odor space) based on two models described above – the order code and primacy. We benchmarked these two models by analyzing cross-concentration and do you cite here cross-individual odor identity predictor. Our analysis shows that both models provide accurate definitions for odor identity which generalize well both across different concentrations within the same animal and across individual animals. The framework also allowed us to test evolutionary predictions of the primacy coding hypothesis [27].

## RESULTS

### Receptor affinity space

To illustrate the odor coding based on the order of OR recruitment, we will use a simplified model of the olfactory system that encodes only two odorants and their mixtures (**Fig. 1**). Based on the OR responses to a particular set of odors (X and Y, **Fig. 1A,B**), ORs can be represented as points in the 2D space with coordinates describing their affinity to the odorants (**Fig. 1C**). Specifically, ORs with higher affinity to an odor have larger coordinates representing that odor, while those with lower affinities have smaller coordinates (**Fig. 1D**).

**Figure 1.**
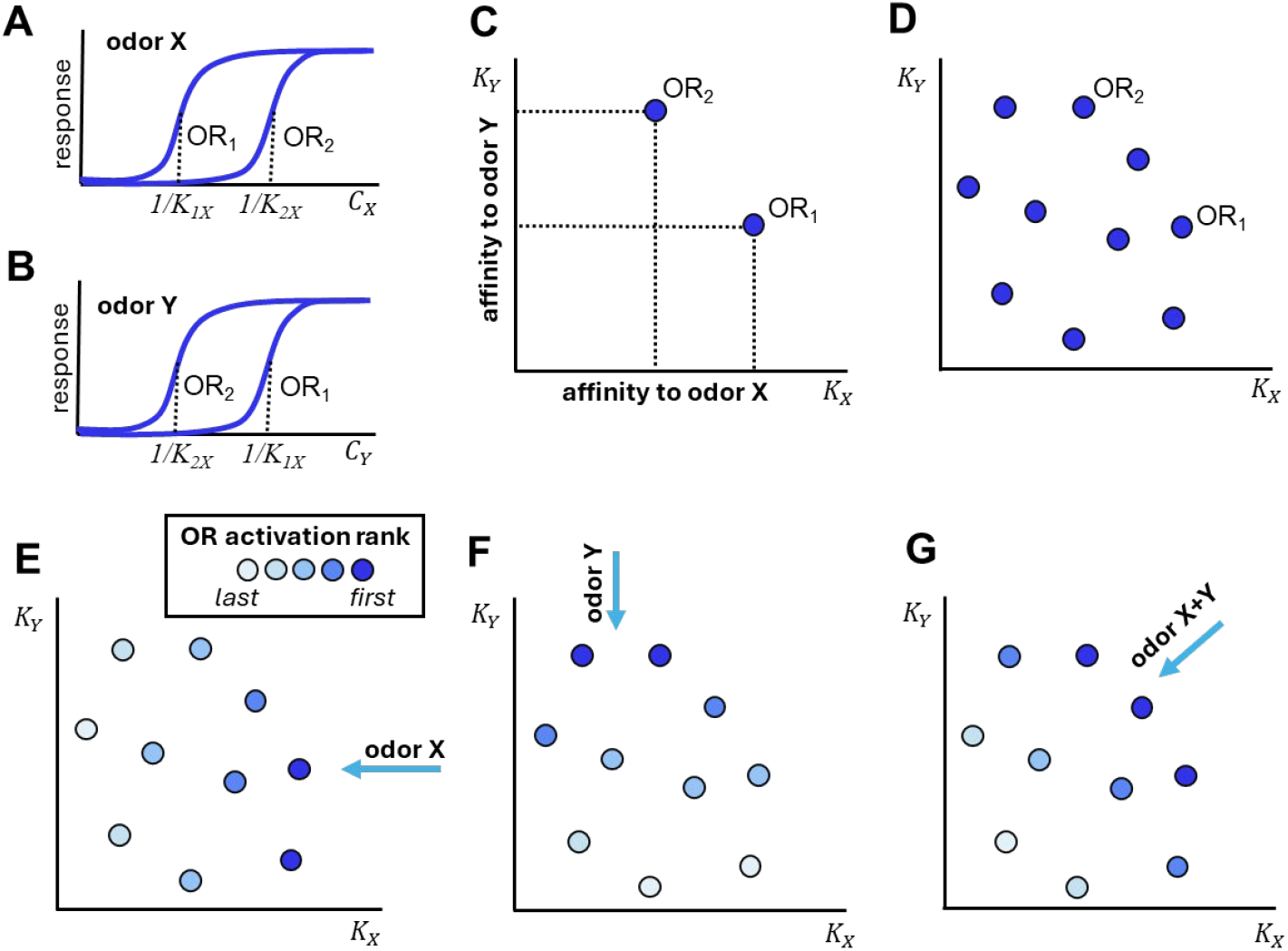
Odorants are represented by waves of OR activation propagating in the receptor affinity space. (**A, B**) An illustration of dose response curves for two olfactory receptors, OR1 and OR2, in response to two odors X and Y as a function of concentration. OR affinities are denoted by *K*_*X*_ and *K*_*Y*_ and can be found from the concentrations at half activation, e.g. *K*_*X*_*=1/C*_*50*_. (**C**) A simplified 2D receptor affinity space that corresponds to the two odors. OR1 and OR2 are arranged according to their affinity to X and Y. (**D**) More ORs included in the space. (**E**) The order of OR activation in response to odor X is determined by their arrangement along the corresponding affinity axis. Odor X is represented by a vector that corresponds to the direction of receptor activation. (**F**) The activation of ORs in response to odor Y moves along axis Y. (**G**) A mixture of two odors X and Y in equal parts activates receptors along the diagonal direction.

If odor concentration increases gradually, odor presentation evokes temporal activation waves propagating in the affinity space. Indeed, as the concentration of odor X increases within a sniff, and assuming quasistationary receptor activation, ORs with the highest affinity to that odor (rightmost in **Fig. 1E**) are activated first, followed by those with lower affinity. This produces an orderly activation wave that propagates through the affinity space from right to left in response to the presentation of odor X. Similarly, odor Y presentation leads to the activation wave propagating from top to bottom (**Fig. 1F**). For mixtures of two odors, X and Y, the activation wave moves through the affinity space in a direction that reflects how much of each odor is present in the mixture (**Fig. 1G**). If the mixture contains more of odor X, the wave is oriented closer to the direction associated with X; if it contains more of odor Y, it tilts toward Y (see Ref. [27] for a more formal model). Overall, the direction of OR activation waves (blue arrows in **Fig. 1E-G**) provides an odor representation defined by the sequence in which ORs are activated.

With more than two molecules present, the affinity space may become high-dimensional, with one dimension per every odor. Its intrinsic dimensionality, however, has been hypothesized to be much lower [28–30]. This reduction occurs because OR affinities are correlated -- many ORs respond similarly to overlapping molecular features, making the effective space a low dimensional. The intuition developed by the simple low-dimensional model presented here becomes relevant. In the following sections, we applied this framework to construct the OR affinity space using glomerular activity data obtained using calcium imaging. We estimated the dimensionality of the affinity space to confirm whether it is low-dimensional. We then verified that odors can be described as activation waves within this space. Finally, we tested the odor space by building a cross-individual and cross-concentration odor identity predictor.

### Imaging glomerular responses to odorants

To construct the receptor-odor affinity space, we imaged glomerular responses to large sets of odorants in several mice (**Fig. 2A**). We obtained spatiotemporal glomerular patterns in response to odor in multiple mice with chronically implanted windows over the dorsal OB, across multiple imaging sessions for odorant presented at one or two air dilutions, depending on the dataset (**Table 1**). Overall, we collected three datasets allowing us to test different features of the olfactory code. The first dataset includes glomerular responses to 65 monomolecular odorants measured at two concentrations in 2 mice. This dataset is used for odor space mapping and primacy model testing (odor space mapping dataset, OSM). The second dataset consists of glomerular responses to 24 odorants, both monomolecular and their binary mixtures, in 7 animals. This data is used for the comparison of odor spaces across animals (cross-animal dataset, CRA). Lastly, in a single animal, we explored the representation of mixtures in more detail. We imaged glomerular responses to binary mixtures of gradually varying composition, i.e., when the proportion of each odorant in the mixture changed by 25% at each step. The third dataset consists of responses to 110 odorants, including 53 monomolecular and 57 mixtures (morphing dataset, MOR). We used it to map the odor space and to test the representations of odor mixtures.

**Table 1:**

|  |  | Mono-molecular odorants | binary mixtures | Concentrations | mice |
| --- | --- | --- | --- | --- | --- |
| Odor space mapping dataset | OSM | 65 | 0 | 2 | 2 |
| Cross animal dataset | CRA | 8 | 16 | 2 | 7 |
| Morphing dataset | MOR | 53 | 57 | 1 | 1 |

**Figure 2.**
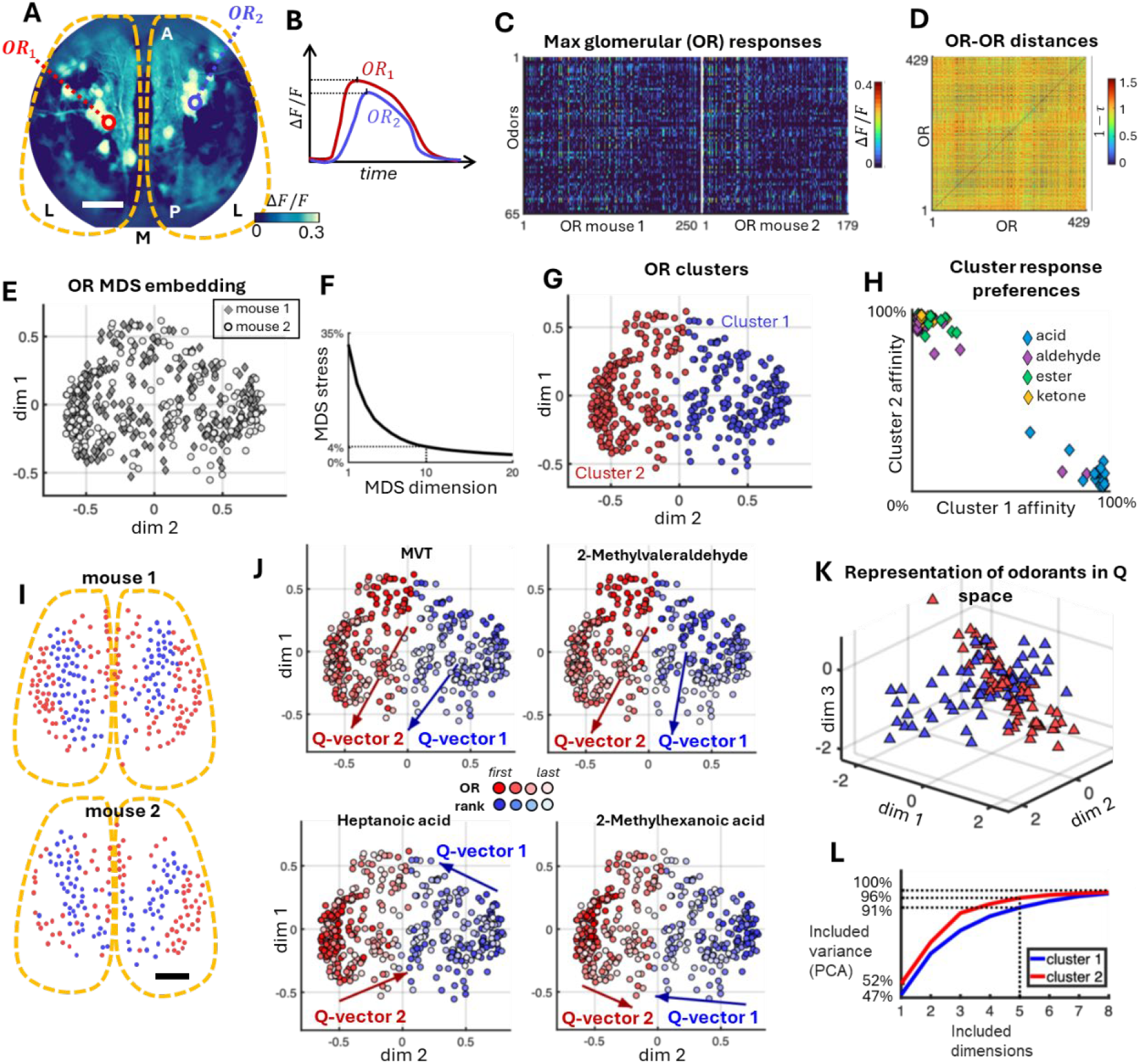
Odor affinity space in the mouse OB. (**A**) An example of trial-average GCaMP6f image of mouse dorsal OB activation during odor presentation (average dF/F over 40-200 msec after sniff onset, OB outline is illustrated by the yellow line). Two example glomeruli are highlighted with circles. (**B**) Illustration of the two glomerular activation traces from **A** in response to an odor. The response maxima are shown for each odor by the horizontal lines. (**C**) The maxima of glomerular responses during the first sniff cycle after odor onset for 65 odorants and 429 glomeruli in two mice (N_mouse 1_ = 250 and N_mouse 2_ = 179) averaged over trials. (**D**) The pairwise glomerular/OR response similarities defined as Kendall’s tau correlations of maximum responses in **C** computed over odors. Similarities are computed across all glomeruli in two mice. (**E**) 2D projection of the 10D MDS embedding of ORs from two mice obtained using the similarity matrix in **D**. The circles and diamonds denote glomeruli of mouse 1 and mouse 2 respectively (N_total_ = 429). (**F**) MDS stress as a function of the embedding dimension. 10D MDS space includes all but 4% of OR-OR distance data. (**G**) Two clusters of receptors discovered by the density-based watershed algorithm. (**H**) Preferred responses of receptors in cluster 1 and cluster 2 to different chemical groups of odorants. Every point’s position was randomly offset by a small amount for better visualization. (**I**) Physical locations of cluster 1 and 2 ORs on the surface of the OB. The OB outline is shown schematically by the yellow dashed lines. (**J**) Examples of temporal activation of ORs for four different odors, visualized independently for the two clusters using color. The directions of OR activation waves are determined by Q-vectors [Equation (1)] shown for each cluster. (**K**) Q-vectors for all odors in the dataset plotted for each cluster (red and blue markers for clusters 1 and 2 respectively). (**L**) Q-space for cluster 2 is somewhat lower dimensional than that of cluster 1 as determined by the principal component analysis (PCA). The scale bars in **A** and **L** are 0.5mm.

### Odor affinity space construction

Using Ca imaging data, we determined the relative locations of ORs within the affinity space. The position of each OR is determined by its affinities to the odorants as described above (**Fig. 1**). Our data, however, does not explicitly contain receptor affinities; we need to use its proxy to position receptors in the affinity space. We reasoned that receptor affinity to an odorant is correlated with the maximum response (response amplitude) of this receptor to the odorant (**Fig. 2B, C**). Subsequently, the similarity between two glomeruli was defined as Kendall’s tau rank correlation coefficient *τ* between their response amplitudes over the set of odorants [26, 31]. Using the similarities, we computed the pairwise distances between receptors in the affinity space - the bigger the similarity, the smaller the distance (*D* = 1 - *τ*, see Methods, **Fig. 2D**). Based on OR-OR distances, and our assumption that higher affinities yield stronger response, we can evaluate the relative positions of receptors in the affinity space by applying the Multidimensional Scaling (MDS) algorithm [32, 33].

Using this procedure we embedded the OSM dataset, which contains glomerular/OR responses in two mice to the same set of odorants, into a shared affinity space (**Fig. 2E**). The dimensionality of the resulting embedding can be evaluated using MDS stress, which represents the normalized difference between observed and computed OR-OR distances in the MDS space (**Fig. 2F**). For a high-quality embedding, 5% of residual stress is considered acceptable [32, 33]. For the OSM dataset, embedding into a 10D space resulted in an MDS stress of 4%, implying that a 10D embedding preserves about 96% of the original distance relationships, with only small distortions. We conclude that receptor affinities are confined to a relatively low-dimensional space (**Fig. 2F**), as assumed in our model (**Fig. 1**), even when ORs from different animals are included in the same embedding.

### Receptor embedding reveals parallel information channels

Our embedding revealed that the ORs do not sample the receptor affinity space uniformly. Two regions on the opposite sides of the receptor distribution contain local aggregations of ORs (negative and positive values of dim 2 in **Fig. 2E**). What distinguishes these two groups of receptors in terms of their functional properties? To address this question, because there is no sharp boundary between the two groups, we applied unsupervised density-based Watershed algorithm [34] (Methods) to split ORs into two groups: cluster 1 and 2 (Fig. 2E).

To establish whether the clusters prefer specific groups of odors, we focused on the analysis of the earliest-responding glomeruli/ORs (primacy sets). In the presented analysis, we used p=10 earliest activated receptors for each animal. For each odorant, we computed the fraction of its primacy set contained within each cluster. For example, if the earliest activated glomeruli/ORs (primacy set) for an odorant are found only in cluster 1, then f(cluster 1) = 100% and f(cluster 2) = 0%, indicating that cluster 1 is sensitive to that odor type (for example, acids). If an odor activates both clusters equally, then f(cluster 1) = f(cluster 2) = 50%. By comparing f(cluster 1) and f(cluster 2) across odorants, we determined how each cluster is tuned to different odors. (**Fig. 2H**). After separating odorants based on their chemical functional groups, we found that receptors in cluster 1 exhibit a bias toward acids, while cluster 2 receptors show a preference for ketones, aldehydes, and other compounds (**Fig. 2H**). We confirmed that the OR tuning shown in **Fig. 2H** remains qualitatively unchanged when the primacy set size p is varied between 5 and 20. Previous studies have shown that similar preferences for functional groups of odorants are displayed by the class I and class II ORs [35]. This comparison suggests that the clusters that we identified based on the neural responses in the affinity space may represent two OR classes defined phylogenetically, derived from homologies between genetic sequences [36–39].

To further compare the identified clusters with OR classes, we examined the spatial distribution of their glomeruli on the OB surface (**Fig. 2I**). Class I glomeruli are known to form a distinct domain on the dorsal surface of the OB [35], spatially segregated from the region occupied by class II receptors. We found a similar arrangement of cluster 1 and cluster 2 glomeruli within our imaging window (**Fig. 2I**). This correspondence strengthens the putative parallel between the functionally defined glomerular clusters and the genetic OR classes. Overall, our data suggests that cluster 1 and cluster 2 glomeruli may correspond to class I and class II ORs, respectively.

### Odors evoke orderly waves of OR activations in the receptor space

As time within the sniff progresses, more glomeruli are activated by odorants. To analyze the dynamics of glomerular recruitment within the sniff, we first measured latency of glomerular activation after sniff onset. To combine multiple animals, we used a continuous set of latencies for each odorant (Methods). For each glomerulus, we then evaluated its rank of activation in the sniff cycle.

To visualize how the activation spreads in each receptor cluster, we highlighted the receptors based on their activation rank – the earlier/later glomeruli/ORs had more/less color (**Fig. 2J**). This visualization showed that receptors were activated in an ordered sequence, resembling a wave of activation moving through the OR ensemble. This observation is similar to the prediction of our theory (**Fig. 1E-G**). To validate this observation quantitatively, for every odor and every cluster, we defined the direction of activation front propagation as a Q-vector, based on the glomerular/OR activation ranks and their position in the MDS receptor space, **x**_r_:

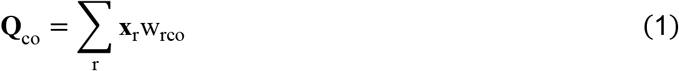

Here w_rco_ are rank-dependent weights computed for each cluster c in response to odor o. For vector **Q**_co_ to represent the direction of activation wave propagation, the earliest receptors are expected to enter this sum with positive weights, while later receptors should have negative weights. This way, Equation (1) can differentiate the coordinates of early and late activated receptors and compute the vector in the direction opposite to the wave propagation. To implement this reasoning, we defined the weights based on the ranks of OR activation combined in the matrix S_rco_ for each receptor r in cluster c in response to odor o, early/later activated ORs have small/large ranks S_rco_. To compute receptor weights in Equation (1), we used the following equation: 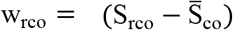, where 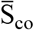 is the S_rco_ averaged across all receptors and the normalization factor Z ensured that the weights have unit length, i.e. ‖**w**_co_‖ = 1. We computed Q-vectors for each OR cluster separately and displayed them in the MDS space (**Fig. 2J**).

Using Q-vectors, we further tested the hypothesis of ordered wave propagation in OR space. For each cluster, we randomly split receptors into two non-overlapping halves. We computed Q-vectors from one half, then projected the positions of the remaining receptors onto these Q-vectors to predict their activation ranks [see Methods, obtaining ranks in the affinity space model]. We found that the predicted ranks for the held out set of receptors appeared to be in high correlation with the observed rank (**Fig. S2)**. This correlation was not present for the randomly shuffled ranks (**Fig. S2**). We thus concluded that odors are represented by robust activation waves that have well-defined directions of propagation (Q-vectors) which generalize across non-overlapping sets of receptors. This result validates visually observed gradients of ranks in **Fig. 2J**.

One possible explanation for the observed activation waves is a correlation between receptor activation latencies and response amplitudes. Indeed, strongly activated ORs are expected to respond earlier in the sniff cycle. Because OR positions in receptor space are determined by their response amplitudes, a strong correlation between latencies and amplitudes could, in principle, account for the emergence of activation waves. To test whether this correlation alone can explain the phenomenon of waves, we performed a control analysis in which both response amplitudes and activation latencies were shuffled using the same random ordering. This shuffling preserved the correlation between amplitudes and latencies. As described above, we then computed Q-vectors representing the activation waves and projected the locations of a held-out set of ORs onto these Q-vectors to assess how well the wave definition generalized across the receptor ensemble. We observed a substantial drop in the correlation between predicted and observed ranks after shuffling (from r = 0.80 to r = 0.27; **Fig. S3**). Therefore, correlations between response amplitudes and latencies, which remain intact under our shuffling procedure, cannot fully explain the observed OR activation waves. A significant component of the rank ordering in receptor space thus arises from the geometric arrangement of the ORs.

We observed that Q-vector pairs for clusters 1 and 2 aligned for certain odors but diverged for others (**Fig. 2J**). To explore this observation, we visualized the Q-vectors for all odors for each cluster (**Fig. 2K**). These sets of Q-vectors appeared to be low-dimensional, with the first five principal (PCA) dimensions of cluster 1 and cluster 2 capturing 91% and 96% of the variance respectively (**Fig. 2L**). When combined in the single space, the Q-spaces were misaligned and spanned distinct dimensions (**Fig. 2K**). This observation further corroborates our conclusion that ORs belonging to the two clusters form two distinct encoding channels which transmit information about different odorant properties.

### The unified odor space

How can the odor representations from the two olfactory channels, clusters 1 and 2, be merged to form a representation of the unified odor identity? An accurate model for combining information from these channels should produce representations that are stable across odor concentrations and consistent between individual animals. To identify the optimal method for merging the two channels, we analyzed the CRA dataset, which includes glomerular responses from seven mice exposed to a set of odorants at two concentrations. We constructed a unified affinity space, identified clusters 1 and 2 of glomeruli, and computed Q-vectors separately for each mouse at each concentration (**Fig. 3A, B**). We then evaluated three alternative methods for combining the Q-vectors from the two clusters to produce a unified representation of odor identity (**Fig. 3C**).

**Figure 3.**
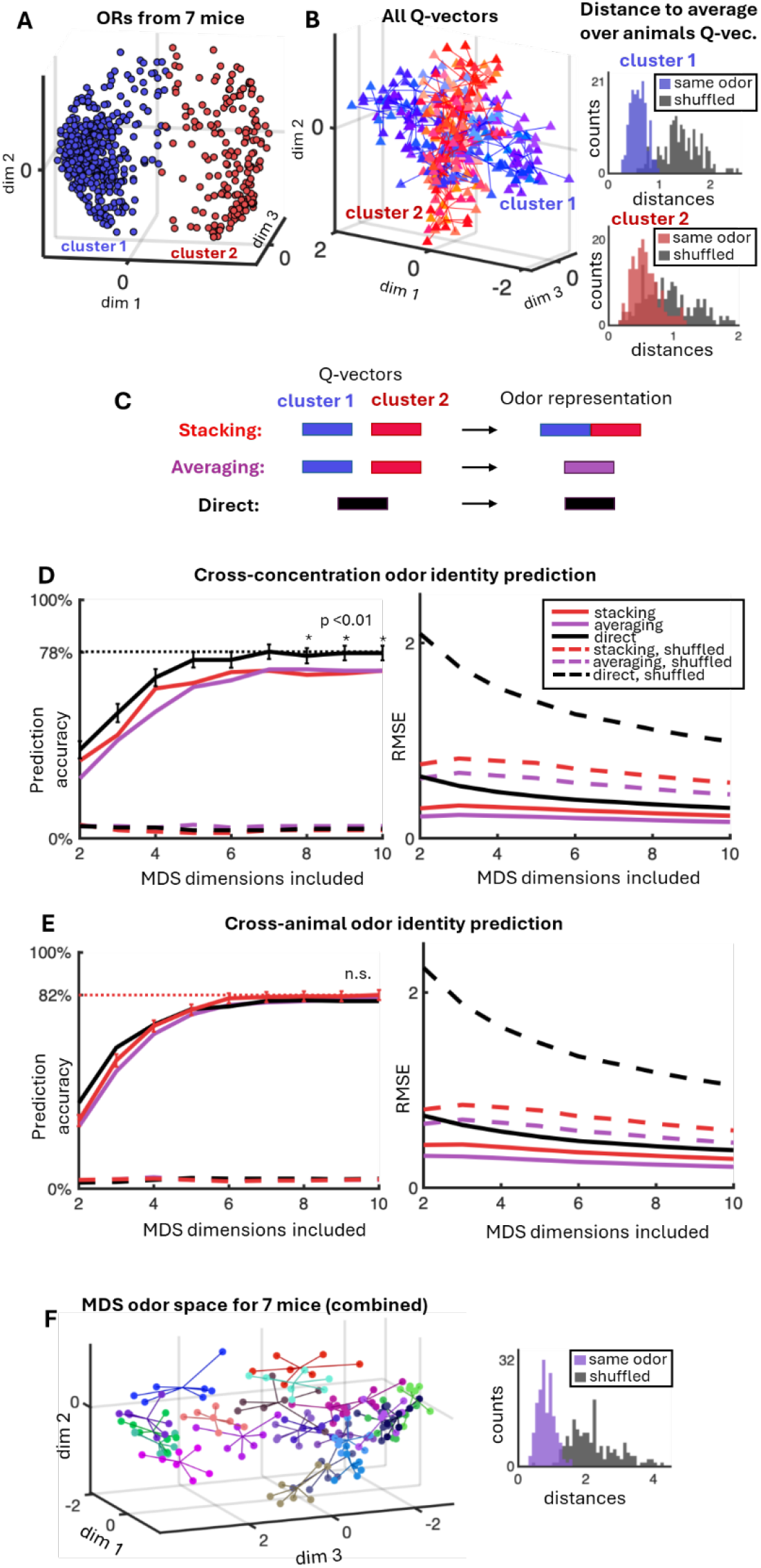
Unified odor space. (**A**) A combined MDS receptor embedding for seven mice using CRA dataset. Two clusters of receptors discovered by the watershed algorithm are shown in blue and red. (**B**) Q-vector spaces of the seven mice. Every point is a Q-vector corresponding to an [odor, mouse, cluster] combination. Q-vectors for the same odor and cluster are linked across mice. The histograms on the right compare the distributions of distances between Q-vectors for the same odor across different mice to those obtained from shuffled pairs. (**C**) Three methods of Q-vector integration into a united odor identity representation. (**D**-**E**) Results of odor identity prediction across animals/concentration as a function of receptor space embedding dimension. (**D**, left) Performance of odor identity prediction across concentration regimes. The performance is shown as a percentage of correct odor identity assignments using leave-one-out cross-validation. The dashed line denotes the maximum achieved performance and colored with a color of the corresponding method of integration across OR clusters. (**D**, right) RMSE measured between odor identity representations corresponding to the same odor presented at different concentrations. (**E**) Performance of odor identity prediction across different animals. The performance is shown as a percentage of correct odor identity assignments (**D**, left) and RMSE measured between odor identity representations corresponding to the same odor in different animals (**D**, right). (**F**) Odor space constructed using the winning “direct” method. The representations of same odor in different animals have the same color and are linked. The distributions of the link lengths (right, purple) is different from random (right, gray).

In the first method, we concatenated Q-vectors **Q**_1o_ and **Q**_2o_ from clusters 1 and 2 for a given odor o to produce the odor representation **Q**_o_. We called this method “stacking” (**Fig. 3C**):

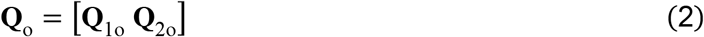

In the second method, “averaging”, we computed the average of the two cluster representations **Q**_1o_ and **Q**_2o_ to obtain **Q**_o_:

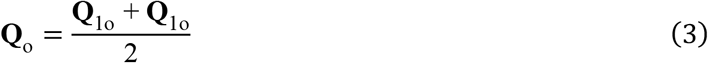

Finally, in the third “direct” method we disregarded the separation of the receptors into clusters and constructed the Q-vectors directly from all receptors across both clusters, using a modified equation of Q-vector computation:

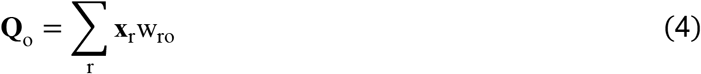

Here **x**_r_ represents one mouse’s MDS receptor coordinates, weighted with rank-dependent weights w_ro_ as in (Equation 1) and w_ro_ is computed for all mouse receptors in response to odor o without separating them into clusters.

We then built cross-animal and cross-concentration odor identity predictors to compare the three integration methods based on their odor identity prediction accuracy. To validate prediction quality, we use leave-one-out cross validation (LOOCV) procedure (Methods) [40, 41]. For the cross-animal predictor, one odorant in one animal was designated as the test data. We computed its combined Q-vector using one of the three integration methods described above. Because glomeruli from all seven mice were embedded in a shared receptor space, the resulting Q-vectors required no additional cross-animal alignment. We then computed the average over animals Q-vectors for each odorant across the remaining non-test data, yielding templates that served to identify the test odorant. The test odor was classified as the identity of the nearest template in the Q-vector space. This procedure was repeated by designating each odor in every animal as testing one-by-one, yielding the success rate for cross-animal odor identity prediction. A similar LOOCV procedure was applied across concentrations, designating one odor at one concentration as the test case. The probability of correct identification in both cross-animal and cross-concentration prediction was used to compare different methods of integrating information from the two OR clusters.

All three integration methods (stacking, averaging, and direct) outperformed their respective shuffled controls and achieved ~80% accuracy for both tasks (**Fig. 3**). In the cross-animal prediction task, the three methods performed similarly, with no statistically significant differences (**Fig. 3E**). However, in the cross-concentration task, the direct method outperformed the others (**Fig. 3D**). Overall, the direct method yielded the most competitive performance across tasks. This result suggests that the separation of odor information into two channels, which is ignored by the direct method, may not be exploited by downstream networks for decoding odor identity. Instead, a global OR activation rank, continuous across channels, appears to provide the most accurate representation of odor identity.

### Primacy theory for odor identity

Above, we tested the order code model for odor identity in the olfactory bulb. In this model, an odor identity is determined by the sequence of OR activation ranks. Alternatively, the primacy model considers only whether a receptor is activated early in the sniff cycle (a primary receptor with low rank) or later (a non-primary receptor with high rank). The receptor space in the primacy model can be constructed similarly to that in the order code model (**Fig. 4A**). Within this space, primary receptors for each odor are expected to lie along the external, high-sensitivity boundary of the receptor cloud (**Fig. 4A–C**). The union of all primary receptors across odors forms a structure known as the primacy hull [42], while receptors that do not belong to any primacy set are termed null receptors (**Fig. 4D**). One of the hypotheses of the primacy model is that null receptors are removed from the genome (become pseudogenes) through evolution [42] (**Fig. 4E**). Accordingly, the number of null receptors is expected to be low (**Fig. 4F**).

**Figure 4.**
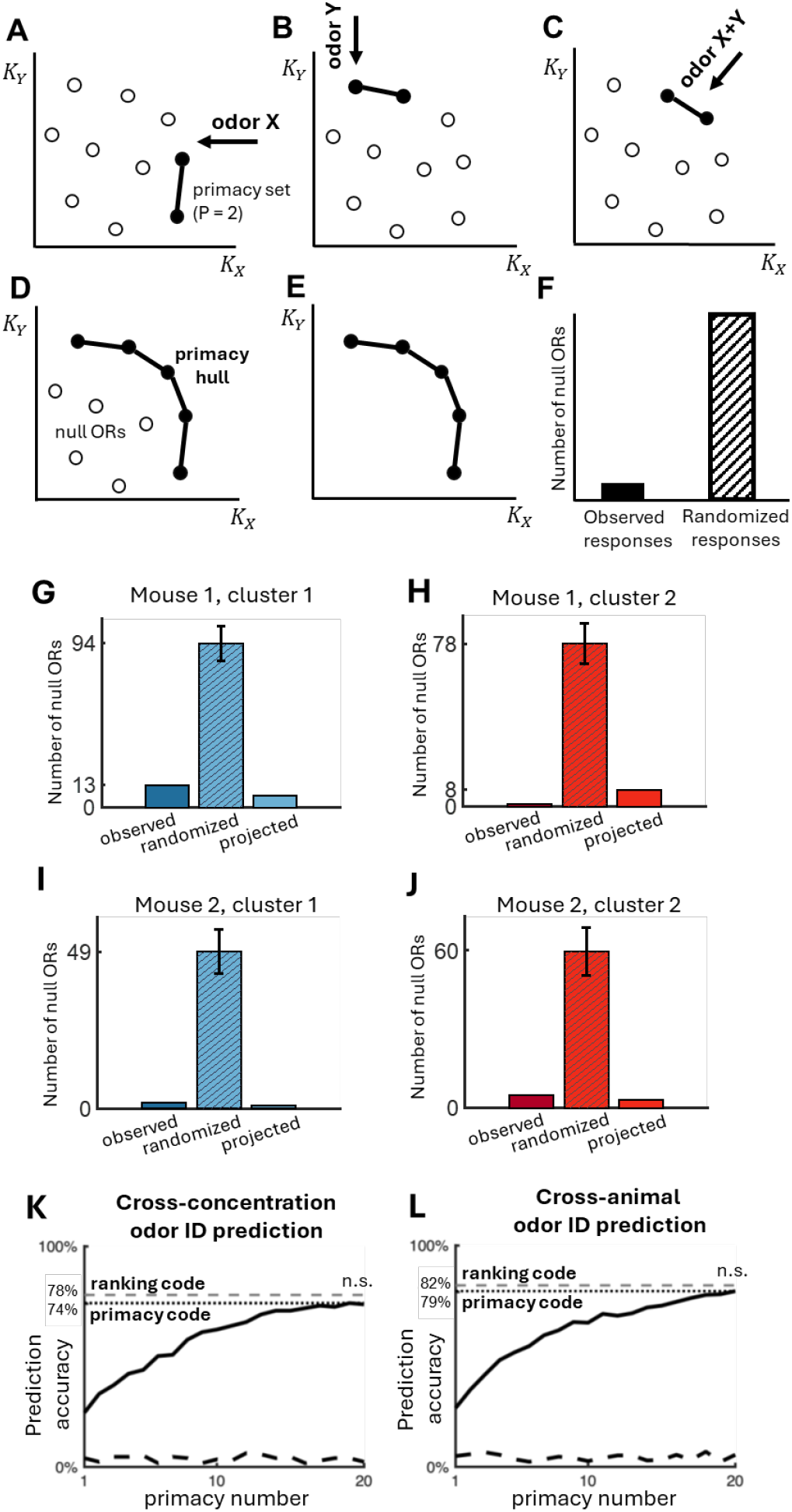
Primacy model for odor identity. (**A**-**C**) Primacy sets formed in response to odor X (**A**), odor Y (**B**), and their mixture (**C**). OR affinities are denoted by K_X_ and K_Y_. The directions of OR activation in response to each odor are shown by arrows. Two earliest receptors (p=2, full circles) are assigned to a primacy set for each odor, with later activated receptors (empty) considered non-primary. (**D**) Primacy hull consists of all ORs that belong to at least one primacy set (full), with other receptors called null (empty circles). (**E**) The primacy model hypothesizes that null ORs are eliminated through evolution. (**F**) The number of null ORs is predicted to be small in the primacy model (left bar). For a random OR distribution in the affinity space, the number of null ORs is expected to be larger (right bar). (**G**-**J**) The number of null ORs calculated within every cluster and every mouse (left bar in every subpanel) using OSM dataset (65 odorants). The numbers of null ORs in the randomized case (middle) and in the non-random case obtained using the Q-vectors (right) are shown for comparison. (**K**-**L**) Performance of odor identity prediction using the primacy model (percentage of correct odor identity assignment with leave-one-out cross-validation) across concentrations (**K**) and across different animals (**L**). The dashed lines show the results produced by the best order coding model (Figure 3) for comparison. The dotted line shows the best result for the primacy model. The differences between the primacy and the order code model are insignificant (n.s.).

Several issues complicate testing this hypothesis directly. First, one cannot measure responses to all possible odorants present in the environment. Therefore, some of the ORs may appear to lack associations with any primacy set because their odor is missing from the experimental set of odorants. Second, one cannot assume that all irrelevant ORs are immediately eliminated. Some ORs may be present in the genome as artifacts or due to their non-olfactory roles, despite their irrelevance to the primacy mechanism. Finally, since the identities of primary receptors for each odorant depend on the value of the primacy number p, a parameter that is not known a priori, the information about the frequency of use for each OR cannot be exactly determined. In view of these complications, we use a statistical strategy that computes the frequency of OR use and compares this frequency to randomized data. The prediction of the primacy theory is that real ORs are used more frequently as members of primacy sets compared to randomized data.

We tested this prediction using the OSM dataset which contains responses to 65 odors for two mice. As we established earlier, receptor space contains two clusters of ORs with different odor tuning properties (**Fig. 2**). We expect that these two clusters form independent primacy sets, and the hypothesis formulated above has to be tested for them separately. We determined the primacy sets for each odor using p=10 in each animal (Methods). This analysis shows that the number of null receptors was low for both clusters and for both animals (**Fig. 4G-J**, first bar in every panel, “observed”).

We then compared this result to randomized data. Our expectation was that a random distribution of receptor affinities would exhibit a higher number of null ORs. To generate such a random set in the receptor space, we preserved the marginal distributions of OR densities projected onto each coordinate axis for each cluster. This randomization procedure displaced ORs from the surface of the OR cloud into its interior, without altering the projected distributions. Practically, this procedure is accomplished by shuffling OR coordinates for each axis/mouse/cluster combination.

To obtain primacy sets for ORs with randomized coordinates, we evaluated the responses of this randomized ensemble to the full set of odorants. To preserve the geometry of the odor space, we used the same Q-vectors as in the non-random case. For each odorant, we computed OR activation ranks by projecting randomized receptor coordinates onto the corresponding Q-vector using the affinity-space model (Methods). These ranks were then used to derive primacy sets for each odor. As predicted by the primacy model, the number of null ORs in the randomized case was significantly higher than in the unperturbed case (**Fig. 4G–J**, second bar in each panel, “randomized”).

So far, we have presented the results suggesting that the number of null ORs based on real unperturbed glomerular responses (left bars, **Fig. 4G-J**) is significantly lower than in the randomized case (central bars, **Fig. 4G-J**, p < 0.001, bootstrap). In the unperturbed case, OR activation ranks and primacy sets were derived directly from glomerular responses.

Generating the randomized responses involved two modifications. First, we randomized receptor affinities by shuffling their coordinates in the receptor space. Second, we computed OR ranks for this new set of affinities using the activation model defined by Equation (5). Our goal is to show that the observed differences between the unperturbed and randomized OR sets stem from differences in the OR distribution, rather than from the activation model itself [Equation (5)]. To evaluate the impact of the approximation introduced by the activation model, we applied it to the unperturbed set of OR affinities. The resulting number of null ORs was comparable to that observed in the unperturbed data (**Fig. 4G-J**, first vs. third bars in each panel, “observed” vs. “projected”). This result suggests that the elevated number of null ORs in the randomized case arises from the randomization of OR affinities, not from the activation model. Taken together, these findings support the primacy model’s prediction that ORs are distributed in affinity space around a high-sensitivity surface, termed the primacy hull, and that nearly every OR belongs to the primacy set of at least one odorant.

The result that not primary, or null, receptors are statistically less represented in the whole receptor repertoire, suggest that Primacy coding may be a good candidate for odor idientity representation. Thus, we we compared the performance of odor identity prediction using the primacy model versus the order code.

We used the same cross-animal and cross-concentration prediction tasks as shown in **Fig. 3**. For the order code baseline, we used the best-performing method for each task (**Fig. 3**). For the primacy model, we evaluated three methods of integrating odor representations across the two OR clusters (as described above) and selected the best-performing approach (“direct” method, [Equation (4)]). We found that the highest performance achieved was not statistically different between the two coding models (**Fig. 4K–L**). This result suggests that either primacy or order code model can represent odor identity with similar precision. Our findings suggest that, despite using less information than the order code, the primacy model produces odor representations that are equally robust and generalize well across animals and concentrations.

### Odor mixtures lay on the geodesic lines connecting their components

How do odor mixture representations relate to their components in Q-space? One of our datasets (MOR) includes glomerular responses to binary mixtures (A/B) at varying ratios (100/0, 75/25, 50/50, 25/75, 0/100 percent), allowing us to examine the trajectories these mixtures form in odor space as their composition shifts between two pure components. To analyze this, we embedded the receptors imaged in the MOR dataset, identified two OR clusters using the Watershed algorithm (**Fig. S4**), computed Q-vectors for each cluster (**Fig. 5A**), and generated unified odor vectors via the most accurate “direct” method described previously (**Fig. 5B**, Equation (4)). The resulting odor representations were plotted in the odor space, with each set of five dilution points connected by a line to visualize their trajectory (**Fig. 5B, C**). We found that binary mixtures lie approximately along straight lines connecting their component odors, a pattern evident in both individual OR clusters (**Fig. 5A**) and in the unified representations (**Fig. 5B, C**).

**Figure 5.**
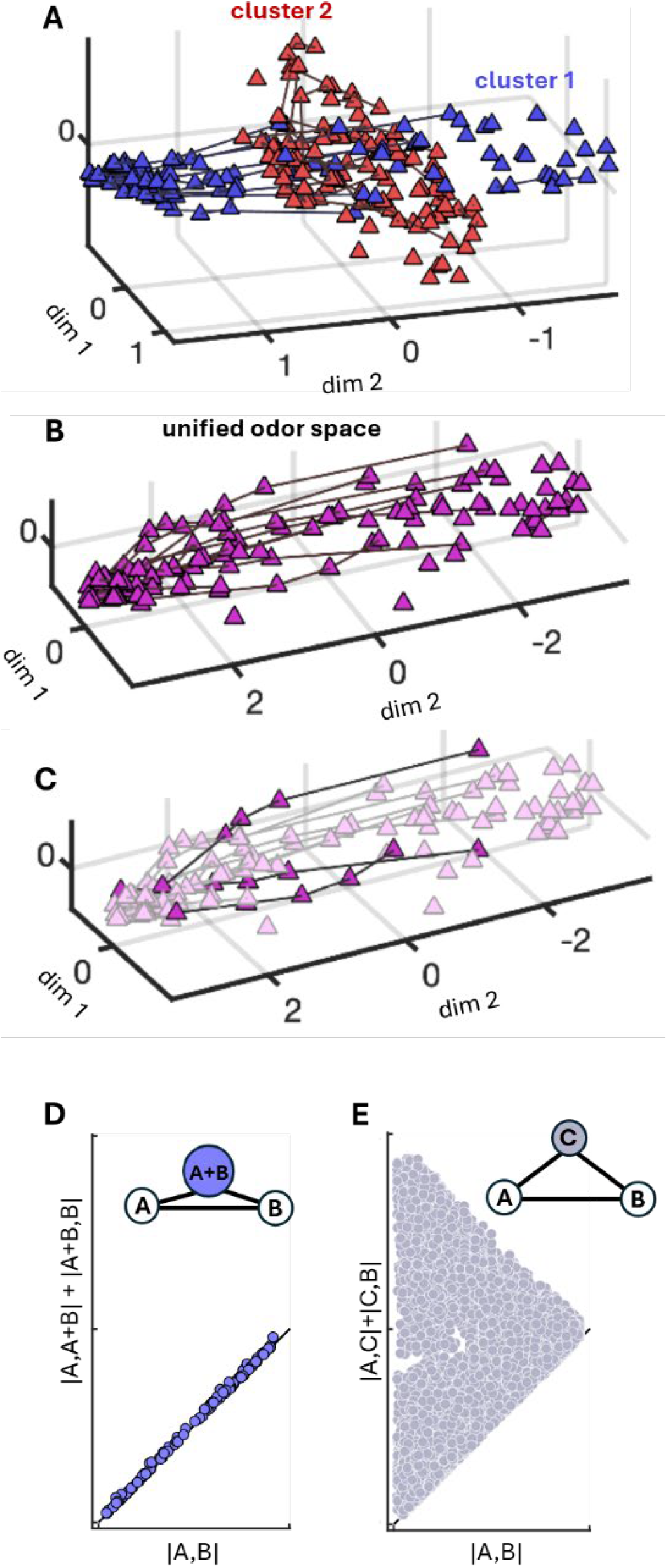
Odor mixtures lay on the geodesics connecting their components. (**A**) Q-vectors based on the clusters 1 and 2 receptors of the MOR dataset. Q-vectors for the binary mixtures of the same components at different concentrations are connected by the black lines. (**B**) The representation of all Q-vectors in the dataset (odor space) unified using “direct” method of odor identity representation. Representations for binary mixtures of the same components are connected by lines. (**C**) The same unified representation as in (**B**) with three mixture series highlighted for visibility. (**D**) Triangle inequality for representations of odor mixtures. The direct distance between a mixture’s components in the odor space is plotted against sum of the distances from the mixture to its components. Every point represents an individual mixture. (**E**) Triangle inequality for every three arbitrary odors in the odor space. Every point is a unique triple of odors A, B, and C.

To quantitatively test this observation, we evaluated the triangle inequality for binary mixtures and their components. Representations of a mixture A+B and its pure components A and B form a triangle in odor space. According to the triangle inequality, the direct distance between A and B should be less than or equal to the sum of the distances from A to A+B and from A+B to B: |A, B| ≤ |A, A+B| + |A+B, B|. If the representation of the mixture lies exactly on the line connecting A and B, this inequality becomes an equality: |A, B| = |A, A+B| + |A+B, B|. We tested whether mixtures lie on straight (geodesic) lines connecting their components by plotting the right-hand side of the equation against its left-hand side. We found that the triangle equality held accurately for all mixtures in our dataset (71 mixtures of binary components; **Fig. 5D**). In contrast, triangles formed by arbitrary odorant triplets (A, B, and C) satisfied the triangle inequality, but with the sum of two sides significantly exceeding the direct distance: |A, C| + |C, B| > |A, B| (**Fig. 5E**). This result indicates that the triangle equality observed for mixtures is not a generic feature of the data. Instead, it supports the conclusion that odor mixture representations lie on geodesic paths connecting their components [43].

## DISCUSSION

The mechanisms by which the olfactory system encodes odor identity are poorly understood. In this study, we collected glomerular responses from the mouse olfactory bulb to large panels of odorants, presented at multiple concentrations in several animals. We tested two models of coding for odor identity in the early olfactory system: the order code and primacy models. Both rely on the concept of a receptor space, where olfactory receptors are organized according to their affinities for odorants. Our analysis of OR responses revealed two clusters of receptors with differing odor tuning properties. These clusters approximately correspond to the two classes of ORs found in the genome [39]. We observed that odorants elicit activation waves that propagate through the receptor space. This observation helped us define odor representations in the order coding model. The directions of these waves, called Q-vectors, differ between the two OR clusters. Q-vectors serve as robust odor identity representations in the order code model, generalizing across both concentrations and individual animals without requiring additional alignment.

We next studied the alternative model (primacy) for its ability to produce concentration- and subject-invariant odor representations. Using leave-one-out cross-validation on cross-concentration and cross-animal prediction tasks, we found that the accuracy of the primacy model was statistically indistinguishable from that of the order code model. Together, these findings suggest that both the order code and primacy models generate equally accurate odor representations that are invariant to concentration changes and generalize robustly across individuals.

A similar combinatorial approach for representing odor identity was used in the studies of insect olfactory system [31] to compare odor-evoked glomerular activity patterns. The approach allows direct comparison of glomerular responses due to exact mapping of glomeruli in the antennal lobe across individuals. Leveraging this anatomical consistency, the authors applied Kendall’s tau rank-correlation [26, 31] to the full glomerular response profiles, assessing the relative ordering of activation across odors. They demonstrated that this rank-based similarity measure outperformed both latency- and amplitude-based metrics in capturing odor identity, with robustness across concentrations and individual animals. In mice, such comparisons are more challenging due to variability in glomerular spatial positioning across individuals. Our approach using Q-vectors made it possible to avoid explicit alignment of glomerular identity and odor spaces, allowing direct comparison of odor identity representations across individuals.

The primacy model encodes odorants based on the rankings of OR responses but simplifies this representation by dividing receptors into two groups: *primary* and *non-primary*. Primary receptors are those that respond earliest to an odorant and can be used to define its identity. This transformation, from a full response rank to a binary primary/non-primary classification, entails a loss of information. Thus, the primacy code relies on less information than the order code to represent odor identity. Remarkably, we find that the primacy model achieves comparable accuracy to the order code in predicting odor identity across concentrations and individual animals (~80% correct identification). This result suggests that, despite its reduced informational complexity, the primacy code captures the most relevant features necessary for representing odor identity.

The residual variance of receptor embeddings decreases rapidly with increasing embedding dimension (**Fig. 2G**), indicating that ORs sample a space of low dimensionality. This finding aligns with previous observations that some ORs respond similarly to odorants sharing functional groups, carbon chain lengths, and other molecular features, resulting in correlated responses across odorants [35, 44, 45]. The low dimensionality of receptor space likely reflects a sampling strategy in which the olfactory system encodes only a limited set of odorant properties relevant to an organism’s survival, rather than the full range of possible molecular features. This strategy parallels that of color vision, where the continuous spectrum of light is represented by a small number of color receptors. Similarly, taste encodes a limited number of nutrient categories through discrete receptor families. These systems operate through labeled lines, with each receptor type sampling a distinct dimension of the stimulus space. Hippocampal place cells use a different strategy, representing 2D space through overlapping, spatially localized place fields, a form of mixed encoding. The olfactory system appears to combine both approaches, employing a hybrid “mixed-labeled line” strategy to sample the low-dimensional odor space. We identified two clusters of ORs with distinct tuning properties. These clusters form independent odor representations (**Fig. 2J**), resembling labeled lines akin to those in the taste system. Within each cluster, odor representations (Q-vectors) are generated through a combinatorial code involving multiple glomeruli, exemplifying the mixed encoding strategy. It is possible that additional labeled lines, corresponding to other genetically defined OR families, exist in regions of the olfactory bulb beyond our imaged dorsal surface. Overall, our findings suggest that the olfactory system employs a unique hybrid sampling strategy, integrating elements from both labeled line and mixed coding mechanisms observed in other sensory modalities.

The primacy hypothesis makes a specific prediction for the case in which ORs sample a space of low dimensionality. In this scenario, when receptor affinities are assigned randomly, many ORs end up inside the overall receptor distribution and therefore do not belong to the primacy set of *any* odorant (**Fig. 4D**). We refer to these receptors as null ORs. Since null ORs are not useful for encoding odor identity, and assuming that they serve no alternative function, they are expected to be eliminated from the genome in the course of evolution. We therefore proposed that, as a consequence of the primacy coding and the low dimensionality of receptor space, most functional ORs should lie along a high-affinity manifold we termed the primacy hull (**Fig. 4D**), while the number of null ORs should be reduced (**Fig. 4E**). Based on our current findings, two distinct primacy hulls are likely formed, one for each of the two identified OR clusters. We assessed the number of null ORs for each cluster and compared these to expectations from random affinity assignments (see Methods). In agreement with the primacy hypothesis, we observed that the number of null ORs was significantly reduced in the glomerular response data compared to shuffled affinity ranks (**Fig. 4G–J**). These results provide empirical support for the primacy coding mechanism, as they reveal predicted signatures of this theory in the observed olfactory glomerular responses.

In conclusion, we evaluated two models for encoding concentration-invariant odor identity in the OB: the order code and the primacy model. Both coding strategies exhibited comparable performance in cross-concentration and cross-subject prediction tasks, even though the primacy model contains less information about odorants than the order code. We identified two distinct clusters of glomeruli (or ORs) with different odor tuning properties that form independent odor representations and form separate primacy hulls. For each cluster, we tested a key prediction of the primacy model that every receptor in the OR ensemble participates in the primacy set of at least one odorant. Together, our findings suggest that odor information in the early olfactory system may be conveyed through two parallel coding schemes, the order and primacy codes, each capable of representing odorants with similar accuracy.

## ACKNOWLEDGEMENTS

This work was supported by the National Institutes of Health BRAIN Initiative grant number U19NS112953 and, in part, by National Science Foundation grant PHY-1748958 to the Kavli Institute for Theoretical Physics.

## METHODS

### Glomerular imaging

We imaged glomerular responses to large sets of odorants at different concentrations in several mice. The mice were expressing GCaMP6f5.11 Ca indicator under *Thy1* promoter in mitral and tufted cells and the recordings were conducted over the dorsal OB from the glomerular layer. For each dataset, ROI maps that correspond to individual glomerulus were drawn manually.

For OSM and MOR datasets, we applied the following procedure. For every glomerulus, the corresponding signal (Δ*F* / *F*) was extracted for each odor presentation trial (5 ~ 15 trials) from −200ms to 1900-2000ms relative to inhalation. For every mouse, odor and concentration, Δ*F* / *F* was averaged across trials, with clean air Δ*F* / *F* subtracted from the averaged result for every glomerulus.

### Receptor responses processing

#### OSM dataset

The dataset includes two mice glomerular activity in response to 65 odors at two concentrations.

For each mouse, odor and glomerulus, Δ*F* / *F* was averaged across two concentration regimes. To make glomerular signal comparable across mice, for each mouse and glomerulus, Δ*F* / *F* was centered across odors. To calculate the odor response latency for each glomerulus, we calculated the standard deviation of its Δ*F* / *F* during −100ms to 0ms relative to inhalation – *σ*. The latency was then assigned to time from 0 to 400 msec from the inhalation onset when the glomerular Δ*F* / *F* exceeded 2*σ*.

#### MOR dataset

The dataset consists of a single mouse glomerular activity imaged in response to 54 monomolecular odorants and 56 binary mixtures. Δ*F* / *F* for a single mouse was used without any further processing. The latencies were obtained as for the OSM dataset.

#### CRA dataset

To limit the effect of light scattering and haemodynamic signal contamination on down-stream analysis, Δ*F* / *F* was cleaned with non-negative matrix factorization (NMF). After spatially aligning Δ*F* / *F* across sessions, each trial was normalized by subtracting the pre-inhalation luminance values. The normalized Δ*F* / *F* data for each mouse was concatenated into a single matrix and decomposed into 300 spatial and temporal factors with NMF [46]. The process of selecting factors to include for reconstructing the denoised data was automated, by calculating the ratio of the inter-/intra-condition variance and retaining the top 50% of factors. The response of every glomerulus was normalized by z-scoring across all trials, and the cumulative sum was calculated over time since inhalation onset for each trial. To obtain the glomerular latency for an odor, the cumulative sum of the response was averaged across the odor trials at the same concentration. The latency was assigned as the minimum between 360 msec and the time when the cumulative sum exceeded its 5*σ*.

### OR affinity space

For every receptor, its extracted maximum responses across odors were ranked from the highest (1) to the lowest (N_rec_, number of recorded glomeruli) with respect to the response magnitude (maximum of Δ*F* / *F* during 0 to 300 ms from the inhalation onset) – from the largest to the smallest. The ranks were then used to calculate Kendall-tau correlation (τ) for each pair of receptors. The pairwise correlation was transformed to pairwise receptor distance as 1 − τ. MATLAB multidimensional scaling (MDS) function *mdscale* was applied to the distances over the range of dimensions D = 1:20 to obtain a receptor embedding satisfying the computed receptor distances. The dimensionality of the embedding was accessed using MDS stress output parameter.

### Receptor space clustering

To cluster the receptors in the MDS embedding space, we imposed a 2D grid onto the receptor distribution (dx=dy=0.1, from the minimum to the maximum of 2D MDS coordinates) and built a 2D histogram containing OR counts in each grid square. We then smoothened the histogram using a Gaussian filter (σ_x_=σ_y_=1.3). We used MATLAB watershed algorithm to separate the receptors into two clusters.

### Computing glomerular ranks from latencies

To find ranks of glomerular activations, we first sorted glomerular activation latencies for each odor/condition in the increasing order. The sorting order represents the recruitment order for the glomeruli. For example, the recruitment order of D, B, A, C indicates that glomerulus D is activated first, B – second, etc. To obtain glomerular activation ranks, the sorting order must be sorted again to yield ranks 3, 2, 4, 1 for glomeruli A, B, C, and D respectively. These ranks were used for the subsequent analysis for evaluating the Q-vectors. When several animals were combined in the same receptors space, a unified continuous latency set was used in evaluating ranks. For latencies reaching the end of our latency window, such as 400 msec for OSM dataset, we assigned random latencies >400 msec to yield randomized high ranks.

### Obtaining ranks in the affinity space model

We can compute the OR activation ranks using the affinity space model. Given a Q-vector **Q**_co_ corresponding to a cluster c and an odor o and coordinates **X**_c_ of OR positions of a cluster c, we inverted the Equation (1) to obtain the vector of rank-dependent weights w_rco_:

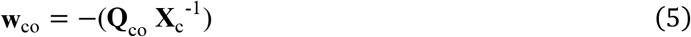

We then computed the receptor activation ranks by sorting the elements of w_rco_, as described in the previous paragraph.

### Randomizing receptor positions in the affinity space

To randomize the receptors in the affinity space, we shuffled every OR coordinate of the matrix **X**_c_. The coordinates were shuffled within each dimension independently on other dimensions, thus leading to the shuffled matrix 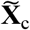. This procedure preserved the marginal distributions of OR coordinates while disrupting potential structure in the affinity space, such as the primacy hull. The shuffled 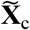 was used in Equation (5) to obtain randomized receptor ranks and the primacy sets as described below.

### Odor identity prediction task

To predict odor identity across concentration regimes for one mouse, we used sets of odor representations that correspond to low and high odor concentrations for each odor 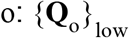 and 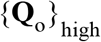, obtained from the two clusters’ Q-vectors using Equations (2)–(4). Then, for each **Q**_o_ in low concentration, we identified a closest vector in 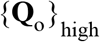 in terms of Euclidian distance. If the identity of the closest representation coincided with o, we marked it as a match, otherwise – as a non-match. We repeated the procedure for each odor in 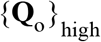 by finding the closest representation in the low concentration odor set. Total number of matches after the two iterations was divided by twice the number of odors to obtain the percentage of matches. The result was averaged across animals. Every method [Equations (2)–(4)] was evaluated independently.

To predict odor identity across individuals, we again evaluated sets of odor representations obtained with Equations (2)–(4) – 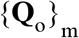 for each mouse m. For a given mouse m and each odor o, we calculated a centroid of all representations of odor o in all other mice ≠ m. Then, in the mouse m, for each odor o, we identified an odor with the closest centroid-representation in other animals. If the identity of the closest representation coincided with o, we marked it as a match, otherwise – as a non-match. We repeated this procedure for all odors and all mice, counting the number of matches. The number of matches was divided by the number of odors and averaged across animals. Each method [Equations (2)–(4)] was evaluated independently.

### Primacy set definition

We identify a primacy set for an odor as p glomeruli with the smallest latency of activation after the sniff onset following by the odor presentation. For Fig. 2H, we used the number of primary glomeruli p=10. We verified that the conclusions about OR tuning in Fig. 2H are not affected if p is within the range from 5 to 20.

### Primacy Q-vector definition

To predict odor identities across animals and concentrations using the primacy model (Fig. 4K, L), and to compare its predictive power to the order code, we defined Q-vectors in this model. We calculated Q-vectors using Equation (1) with a set of weights derived from primacy. In particular, we set 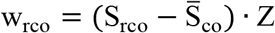, where S_rco_ are 1 for the earliest p receptors r in cluster c in response to odor o, and 0 otherwise. We varied the primacy number between 1 and 20. Here, the normalization factor Z ensured that the weights have unit length, i.e. ‖**w**_co_‖ = 1.

