## Supplementary Figures 1 for "Order code in the olfactory system"

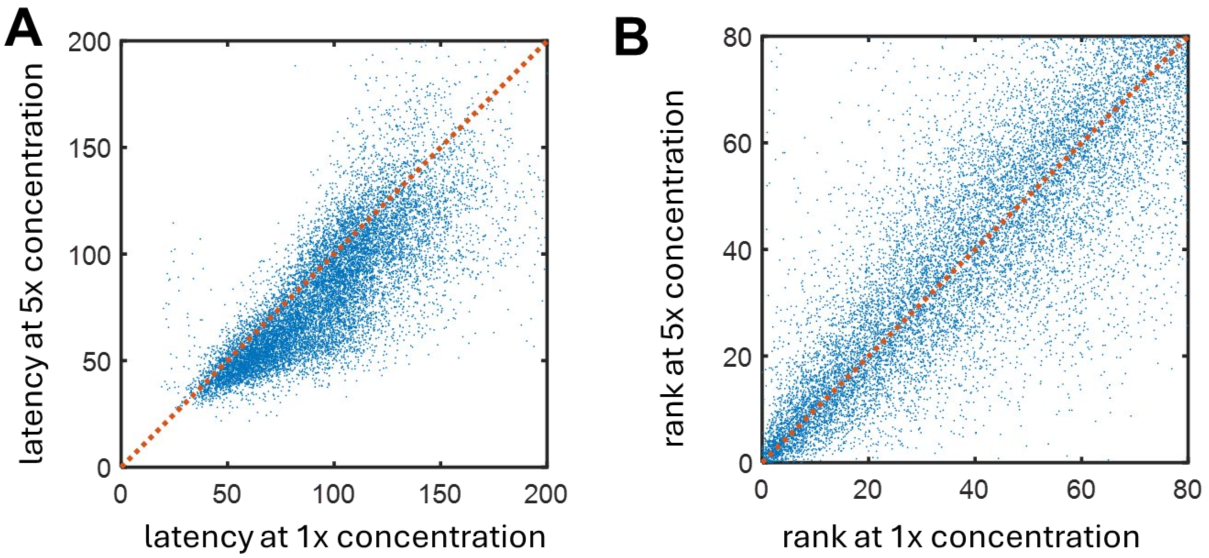


**Figure S1. Glomerular activation ranks but not timing are invariant to concentration changes.** (**A**) glomerular response latencies for all glomeruli-odors pairs in the MOR dataset. The higher concentration latencies are plotted against the lower concentration latencies. That the points lye below the dotted red line indicates that higher concentration latencies are systematically smaller, making latency-based code sensitive to concentration. (**B**) Glomerular activation ranks are approximately concentration-invariant.

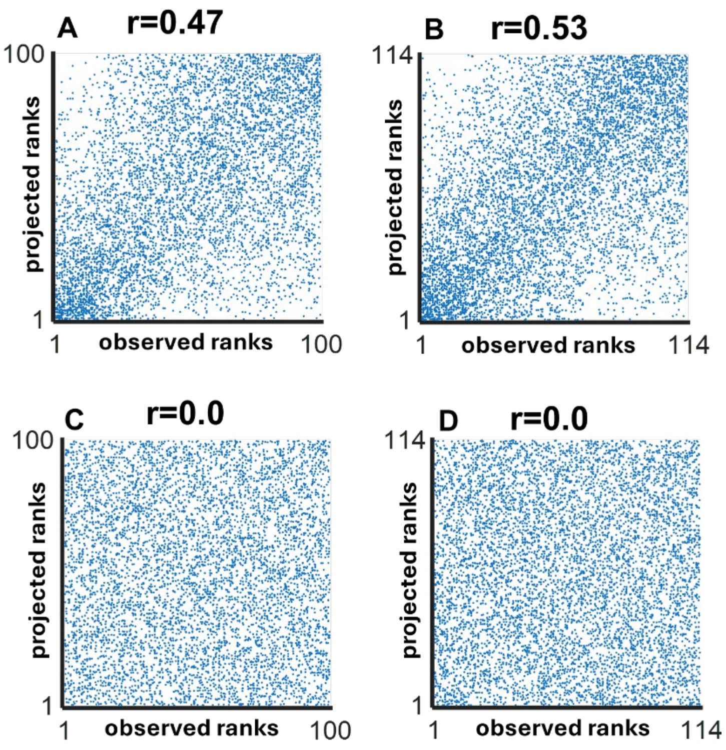


**Figure S2. Activation waves in the OR space are consistent**

**(A-B)** Predicted activation ranks obtained from Q-vectors for the held-out receptors correlated well with their observed ranks. **(C-D)** Similarly obtained predictions using shuffled latencies showed no such correlation.

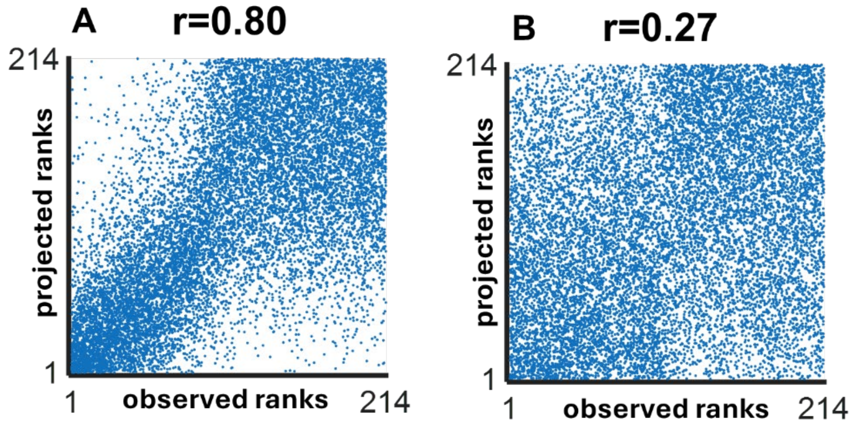


**Figure S3. Observed OR activation waves cannot be explained by the correlation between response amplitudes and latencies.**

**(A)** Predicted activation ranks obtained from Q-vectors for the held-out receptors correlated well with their observed ranks (Pearson r=0.8). This plot is similar to Fig. S2, with the exception that we do not split the receptors into two clusters. This level of correlation suggest that the direction of wave propagation (Q-vector) generalizes well across the receptors.

(**B**) We shuffled OR activation latencies and response maxima using the same random ordering. This shuffling procedure does not change the correlation between activation latencies and response amplitudes. At the same time, the predictability of the OR activation ranks quantified by the Pearson correlation between the observed and predicted OR ranks for the held-out set of receptors is reduced significantly (from r=0.8 in A to r=0.27 in B). This observation suggest that the waves of receptor activation, for the most part, follow from the organization of ORs in the receptor space rather than from the correlation between activation ranks and their response amplitudes.

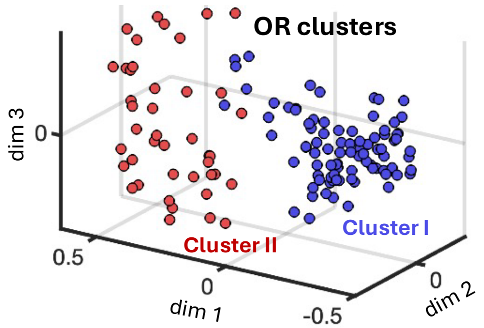


**Figure S4. Receptor clusters based on morphing data (MOR)**
3D projection of the receptor MDS embedding obtained for the morphing dataset. Two OR clusters discovered by the watershed algorithm.
